# Genital microbiota in infertile couples

**DOI:** 10.1101/2023.06.14.544778

**Authors:** David Baud, Adriana Peric, A. Vidal, JM. Weiss, Philipp Engel, Sudip Das, Milos Stojanov

**Author notes:** Equal contribution (David Baud, Adriana Peric). These authors jointly supervised this work (Sudip Das, Milos Stojanov).

## Abstract

Bacteria colonise most of the human body and the genital tract is not an exception. While it has been known for decades that a vaginal microbiota exists, other genital sites have traditionally been viewed as sterile environments, with bacterial presence associated only with pathological conditions. However, recent studies identified specific patterns of bacterial colonisation in most genital sites. Shifts in the bacterial colonisation of the female genital tract have been linked to impairment of reproduction and adverse pregnancy outcomes, such as preterm birth.

The goal of this project is to understand the association between the genital microbiota of couples seeking assisted procreation aid and the outcome of this treatment. Male and female partners will be studied as a unit (“couple microbiota”) and the interaction between their microbiota will be evaluated.

We have characterised microbial samples coming from vaginal and penile swabs, as well as follicular fluid and semen, using next generation sequencing (16S rRNA profiling). The results were linked to clinical data of the patients included in the study and particularly to the results of the fertility treatment process. With this project, we aim to gain a better understanding of how the male genital microbiota could influence the lower (vagina) and upper (follicular fluid) female genital tracts.

## Introduction

Bacteria and other microorganisms colonise most of the ecological niches present in the human body [1–3]. They are not only passive commensals but have a profound influence on the host’s homeostasis, playing a significant role at multiple levels, including protection against pathogens, maturation of the immune system, metabolic pathways, vitamin synthesis, among many others [1,3,4]. Thus, it is not surprising that a disbalance or dysbiosis of the microbiota has been associated with several adverse outcomes.

The impact of the microbiota on the genital tract is not an exception and an increasing number of studies exploring its role on pregnancy, infertility and adverse outcomes are being carried out [5–7]. Most studies focus on vaginal microbiota, which in a healthy state is dominated by members of the *Lactobacillus* genus [8]. Nevertheless, the colonisation pattern may change over time [9]. During the early stages of life, vaginal bacteria form low abundant and highly diverse communities [10]. During puberty, under the influence of sexual hormones, the vagina undergoes to important physiological changes, including the thickening of the epithelium and an increase in the production of glycogen, a condition that favours the proliferation of lactobacilli. Again, with hormonal changes during menopause, the proportion of lactobacilli diminish, leading to a more diverse and less abundant microbiota [11]. Variation of the composition of the vaginal microbiota may also occur at a shorter time scale and may be influenced by multiple factors. Studies on different human population suggest that the genetic background plays an important role in the stability of colonisation by lactobacilli, probably by the interplay of the host’s immune response [12,13]. Women of Caucasian and Asian origin tend to have a more stable, *i.e. Lactobacillus*-dominated, microbiota compared to women of African and Hispanic origin [12].

Recently, bacterial colonisation of other parts of the female genital tract has been characterised [14]. While previously considered sterile, the existence of specific microbial colonisation profiles has been described for the uterus, fallopian tubes and ovaries [14]. Several studies published in recent years, support the notion that a low-biomass microbiota is present in the uterine cavity and the rest of the upper genital tract, forming a continuum with the lower genital tract [14–16]. Compared to vaginal microbiota, there is an increased bacterial diversity, but there is still no clear evidence of the exact impact of the upper tract microbiota on health and reproductive outcome [14]. Increase in the diversity of vaginal bacteria (mostly anaerobic or facultative anaerobic bacteria) is associated with adverse gynecological and obstetrical outcomes. Bacteria that have been associated with a negative impact include, amongst others, *Chlamydia trachomatis*, *Gardnerella vaginalis*, *Prevotella spp.*, *Sneathia spp*., *Bacteroides spp*., *Mobiluncus spp.* and *Atopobium vaginae* [17–20]. This list is certainly not exhaustive as many other species might be associated with vaginal infections [21]. Women with a dysbiotic vaginal microbiota have an increased risk of developing both gynecological (cancer) and obstetrical (miscarriage, preterm birth) pathologies [17,22]. Therefore, unlike other body sites, increased microbiota diversity is negatively associated with gynecological and obstetrical outcomes [23,24]. However, the pathophysiological mechanisms remain unknown as different combinations of genital bacteria can form the same pathology, as it is the case in bacterial vaginosis.

Other than vaginal colonisation, microbiota of the female upper genital has not been extensively investigated, compared to other body sites. Similarly, the male genital microbiota has been mostly neglected. Initial studies on the colonisation of the male genital tract focused on well-known genital pathogens (*Chlamydia trachomatis*, *Ureaplasma spp.* and *Mycoplasma spp.*, among others) or have relied on classical microbiological methods for culturing bacteria [25–28]. This led to the general view that semen is poorly colonised by bacteria, except in cases of ongoing infections where pathogens directly impair fertility. In recent years, bacterial communities colonising the male genital tract have been characterised using next generation sequencing [29–31]. It became clear that semen is not sterile and that bacteria are found not only in infertile men, but also in normozoospermic men (men with normal sperm parameters according to reference values set by the WHO, which include spermatozoa count, concentration, motility and morphology). Most of the studies have been performed on male partners of infertile couples, due to the availability of semen samples that were used for spermiogram analysis [32–35]. Despite multiple sequencing strategies, analysis pipelines and cohorts, similar community types were characterised, suggesting that a specific seminal microbiota might be necessary for optimal sperm function (or that an inadequate seminal microbiota may interfere negatively with sperm function). Presence of lactobacilli was often associated with good semen parameters, but their exact role remains to be elucidated. On the other hand, many bacterial genera found in semen were previously negatively associated with vaginal health and reports indicate that in some cases, sexual intercourse could lead to disturbed vaginal microbiota [36,37]. As an example, *Prevotella* is one of the major genera encountered in semen and has been previously associated with bacterial vaginosis [17]. Interestingly, an increased abundance of *Prevotella spp.* was also negatively associated with semen parameters such as motility and morphology [29,31]. Other semen-associated genera that may negatively impact female genital tract include, amongst others, genera like *Gardnerella*, *Atopobium*, *Sneathia*, *Dialister* and *Finegoldia* [29,31].

Despite the potential impact of seminal bacteria on human reproduction and dysbiosis of the female genital tract, this field of research remains understudied. In addition to investigating the composition of genital microbiota in infertile couples and its potential impact on infertility, our study also aimed to explore the potential interaction between male and female microbiota. By collecting samples from both male and female partners, we aimed to investigate, whether the microbiota of one partner could influence the composition of the other partner’s microbiota. Due to the potential issues associated with analyses involving low biomass microbiota, like semen and follicular fluid samples, we implemented a series of negative controls and stringent *in silico* elimination of possible contaminants to ensure the accuracy and reliability of our results.

## Materials and methods

### Patients and samples collection

This study comprises samples from couples diagnosed with infertility that were obtained from the Luzern Cantonal Hospital (Switzerland) between October 2018 and July 2020. All patients included in the study gave their written consent and utilization of the samples was approved by the Ethics Committee Northwest and Central Switzerland (EKNZ - REPROLUKS003), according to the Swiss Federal Act on Research involving Human Beings.

Samples were collected using eSwab (Copan Diagnostics, Italy) by direct sampling (vagina and penis glans) or by immersion in the biological fluid (follicular fluid and semen). Negative controls consisted of swabs that were opened in the same rooms in which the medical examination took part or processing of sterile water through the pump used to retrieve oocytes and follicular fluid (table 1).

**Table 1.** Description of the sample types.

| Sample (abbreviation) | Description and sampling |
| --- | --- |
| Vagina (vag) | Vaginal swab |
| Follicular fluid (fol) | Immersion of the swab in the follicular fluid obtained from oocyte retrieval |
| Penis (pen) | Swab of the penis glans (self-sampling) |
| Semen (spe) | Immersion of the swab in semen sample obtained for spermogram analysis |
| Control (ctrl) | Swab exposed to the environment in which sampling was performed |
| Follicular fluid control (fol_ctrl) | Immersion of the swab in the sterile medium that has been passed in the aspiration device used for follicle retrieval |
| Technical control (tech_control) | Sterile water sample that has been processed for DNA extraction |
| Water (water) | Sterile water sample used as negative control for the amplification of the V1-V2 regions of the 16S rRNA |
| Community DNA Standard (comm) | Commercial Microbial Community DNA Standard |

ZymoBIOMICS Microbial Community DNA Standard (Zymo Research Corp., Irvine, CA, United States) was used to evaluate the potential variability of the sequencing results between different runs.

### DNA extraction and quantification of bacterial copy number

Tubes containing swabs were vortexed for 30 sec. A total of 500 μl of the tube medium was used for DNA extraction, which was performed with the QIAamp DNA mini kit (Qiagen AG, Basel, Switzerland). After an initial step of lysis using 0.3 μl Ready-Lyse Lysozyme (Epicentre, Madison, WI, USA) at 37°C with 500 rpm shaking for 60 min, extraction was done according to manufacturer’s specifications. Elution was done with 50 μl molecular biology grade H_2_O.

Quantification of bacterial 16S rRNA copy number was assessed by quantitative RT-PCR using 300 nM of both primers F-tot (5′-GCAGGCCTAACACATGCAAGTC-3′) and R-tot (5′-CTGCTGCCTCCCGTAGGAGT-3′) in the Rotor Gene 6000 thermocycler (Corbett Research, Sydney, Australia) with the iTaq Universal SYBR Green Supermix (Bio-Rad, Reinach, Switzerland).

### Bacterial 16S rRNA amplicon sequencing

The variable regions V1-V2 of the 16S rRNA gene were amplified using custom barcoded primers (F-27/R-338) containing Illumina sequencing adapters, as previously described [38]. PCR was done with the Kapa HiFi PCR kit (KAPA Biosystems, Cape Town, South Africa) using 5 μl of extracted DNA as template and the following cycling conditions: 3 min initial denaturation at 95°C, followed by 30 cycles of 30 s at 98°C, 30 s at 56°C and 1 min 30 s at 72°C, with a final extension of 5 min at 72°C. Four libraries were prepared for a total of 282 samples, including the negative controls.

Prior sequencing, PCR products of each sample were evaluated by agarose gel electrophoresis (supplementary figure 1). Based on gel band intensities, samples were classified as “high” (H, strong band), “low” (L, visible band) and “null” (N, absence of a visible band). For each of the four libraries, 3 pools of samples corresponding gel band intensities were prepared (H, L and N). Each pool was purified and quantified, in order to maximise the number of reads for each sample. This step was necessary to avoid the overrepresentation of reads coming from the samples with high bacterial loads (*i.e.,* most of the vaginal samples and a small proportion of penis samples). Bands of each pool were quantified using a Fragment Analyzer (Advanced Analytical Technologies, Ankeny, IA, USA).

Illumina sequencing was performed at the Lausanne Genomic Technologies Facility (GTF) of the Lausanne University using an Illumina MiSeq instrument in paired-end mode 2 x 250 nt (Illumina, San Diego, CA, United Sates).

### Bacterial community analysis

#### Pre-processing and quality filtering

Demultiplexing of the raw sequencing data was performed with the illumina-utils package [39]. Read processing and phylogenetic sequencing analysis was performed with the DADA2 pipeline using default parameters [40] (Supplementary Data, GitHub pipeline). Read quality control, trimming, dereplication and filtering were also perform using the DADA2 pipeline. Samples with less than 1000 reads post processing were removed from the analysis (Figure 1C, kept vs removed samples).

**Figure 1.**
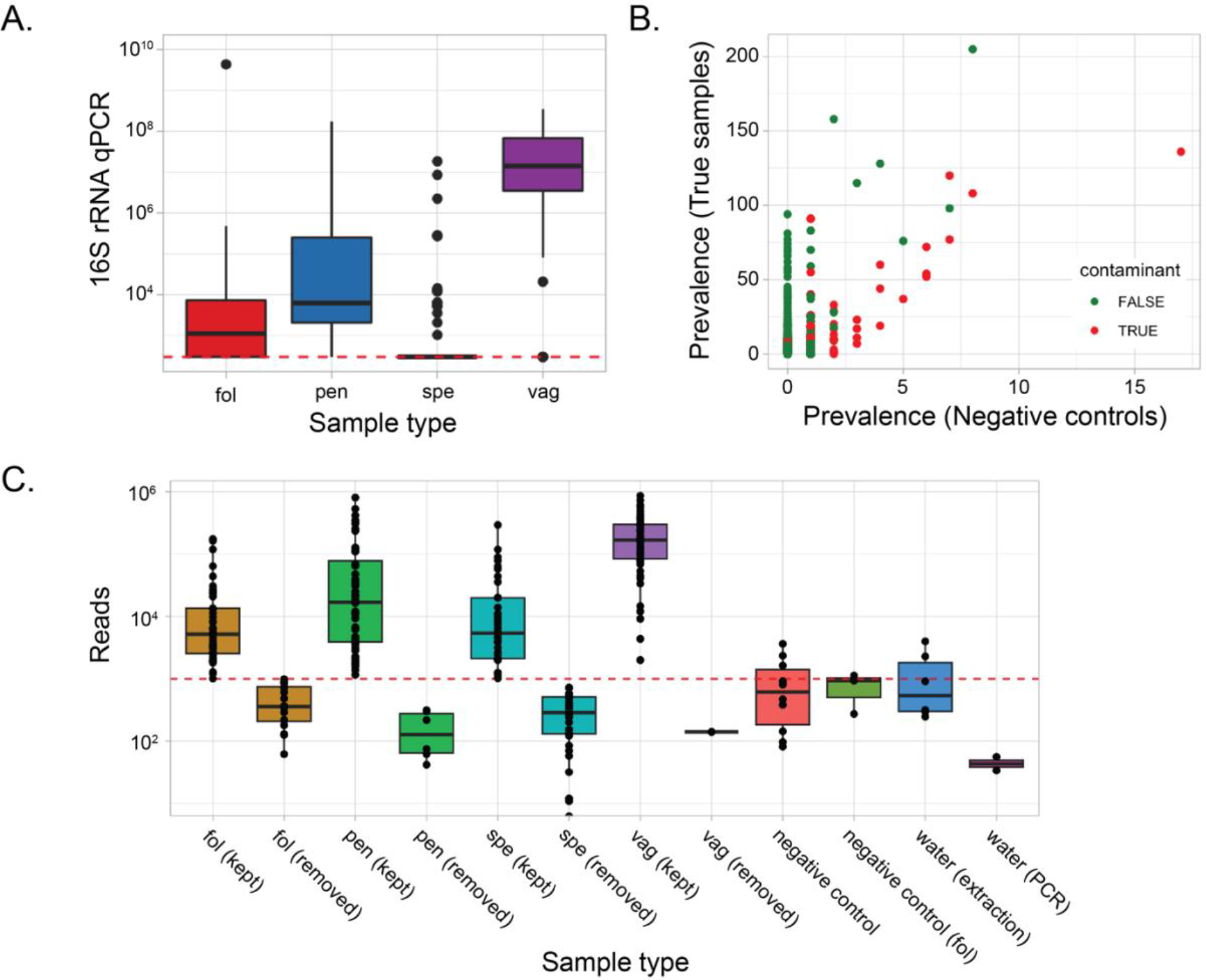
Controlling for low abundance bacterial biomass samples. A. Quantification of bacterial load by qPCR. Partial amplification of the 16S rRNA gene was used to quantify the bacterial biomass in each of the four sample types. The dashed red line represents the limit of detection of 300 copies of the 16S rRNA gene. B. Prevalence of the taxa identified by “decontam” in the true samples and in negative controls. Scatter plot showing the prevalence of ASVs in samples (red) against negative controls (green - extraction and PCR water controls). C. All samples after the filtering steps. Boxplots depicting the reads per sample type, showing samples that were retained in the analysis (kept) and samples that were removed from the analysis (rem). The red dashed line represents the limit of 1000 reads under which samples were removed from the analysis. Fol: follicular fluid; pen: penis; spe: sperm; vag: vagina.

#### Taxonomic assignment and phylogeny

Taxonomy was assigned after alignment with SINA using the SILVA 16S rRNA database (SILVA NR v132) [41], resulting in 16S rRNA gene amplicon sequencing variants (ASVs). Phylogenetic tree was constructed using FastTree version 2.1.10.

#### Decontamination and normalization

Relative abundance of ASVs were converted to estimated absolute abundances by multiplying the 16S rDNA gene copy quantified by qPCR in each sample. Decontamination of sequencing data was performed with the “decontam” package in R [42]. The combined mode, in which both frequency and prevalence methods were used to identify putative contaminants, was used with the default settings.

#### Microbiota diversity analyses

Alpha diversity analyses were performed using the phyloseq function *plot_richness()*, which uses the “vegan” package in R. Results were plotted and analysis of variance (ANOVA) was performed to calculate significant differences between sample types and other metadata variables. Paired sample wilcoxon test was used to investigate differences when categorized into sample type.

To explore the beta diversity the unweighted and weighted UniFrac distances between samples using the *distance()* function in phyloseq was used. Principal coordinate analysis (PCoA) was then applied on the distance matrix using the *ordinate()* function in phyloseq. The resulting PCoA was visualized using the *plot_ordination()* function. Intra- and inter-sample dissimilarities were compared by analysing beta diversity, calculated using the fixed distance method with the “vegan” package. To test the significance of differences, pairwise differences were calculated by permutational multivariate analysis of variance (PERMANOVA) using the *adonis2* function in the “vegan” package.

#### Data visualization and statistical analyses

The majority of statistical analysis and visualizations were performed using Phyloseq, microbiome and vegan R packages were used unless and otherwise specified [43–45]. R package “lefser” was used to infer differentially abundant bacteria [46].

### Data and code availability

All raw sequencing data i.e., fastq files were submitted to Short Read Archive (SRA), National Center for Biotechnology Information (NCBI) under the BioProject PRJNA942221 and BioSample accession numbers SRR23797948 – SRR23798204. All processed data i.e., quality control data, metadata and analysed full datasets are available on zenodo.org with DOI 10.5281/zenodo.7885592 (https://doi.org/10.5281/zenodo.7885592). The full pipeline used for the analysis of sequencing data is available on the following link: https://github.com/dfmemicrobiota/infertile_couples.

## Results

### Study Population of infertile couples

This study comprises samples from 65 infertile couples that were collected at the Luzern Cantonal Hospital, Switzerland between October 2018 and July 2020. We collected 257 samples from four genital tract niches, which consisted of 1) a vaginal swab and 2) follicular fluid from female partners, as well as 3) a semen sample and 4) a penile swab for male partners. Two penis swabs were not available. Infertility reasons were variable, with idiopathic infertility being the most frequently diagnosed condition (Table 1).

**Table 1.** Infertility reason.

| Characteristics | Count (percentage) |
| --- | --- |
| **Age (years)** |  |
| <25 Y | 0 |
| 25-35 | 26 (40%) |
| 36-40 | 27 (43%) |
| >41 | 11 (17%) |
| **Type of sterility** |  |
| Primary sterility | 55 (86%) |
| Secondary sterility | 9 (14%) |
| **Reason for sterility** |  |
| Tubal factor | 5 (8%) |
| Male factor | 16 (25%) |
| Endometriosis I/II | 4 (6%) |
| Endometriosis III/IV | 4 (6%) |
| Anovulation | 4 (6%) |
| Idiopathic | 31 (49%) |
| **Recurrent implantation failure** | 4 (6%) |
| **Previous miscarriage** | 12 (19%) |
| **Recurrent pregnancy loss** | 1 (2%) |

### Bias-control refines genital microbiota analysis and reveals low biomass characteristics

To decipher species composition, we analysed bacterial communities by performing 16S rRNA gene amplicon sequencing and we report bacterial communities based on amplicon sequence variants (ASVs). We found a combined 4001 ASVs in all samples. Considering sample types, 1292 ASVs were found in vaginal swabs, 1111 ASVs in follicular fluids, 2427 ASVs in penis swabs and 1508 ASVs in sperm samples.

Controlling contamination in human microbiota especially in low biomass samples is a critical issue before reporting species structure and composition, due to the possible contamination introduced during DNA extraction and further processing prior to sequencing [47]. We sought to assess this issue by introducing several mitigation steps including absolute quantification of bacteria and inclusion of essential negative controls (n= 18). More specifically, the latter consisted of swabs opened in the room in which sampling was performed (n=8) and washing buffer samples passed through the transvaginal follicle aspiration needle (n=3). In addition, ultrapure water processed by the DNA extraction method (n=5) and ultrapure water used as template for high-throughput sequencing (n=2). First, we quantified sample-wise bacterial copy numbers by amplification of 16S rRNA gene by qPCR (Figure 1A).

Vaginal samples had the highest bacterial load (range: 3.54x10^8^ – 3x10^2^ copies) followed by penis samples (range: 1.75x10^8^ – 3x10^2^ copies). Follicular fluid and sperm samples exhibited the lowest bacterial numbers. Indeed, this reveals that apart from the vagina all other genital tract niches were observed to be of low biomass. Next, we used the statistical method in the “decontam” package in R to infer probable contaminants based on the results obtained with the negative controls (Figure 1B). We investigated read retainment after decontaminations and removed amplicon sequence variants (ASVs) identified as contaminants from the analysis (Figure 1C) for each sample type. Finally, we normalized the data using quantified bacterial copy numbers reporting absolute numbers instead of relative abundance.

### Microbiota analysis reveals insights on the biogeography and gender-specific species composition in the human genital tract

We investigated the species composition and site-specificity of the genital microbiota across gender. We also examined species prevalence and shared bacteria across different sampling sites in each gender. Members of the *Lactobacillus* genus were the most prevalent and abundant bacteria in female samples (both vagina and follicular fluid), with *L. iners* being the most prevalent species, which included two ASVs (ASV1 and ASV11 with 95.2 % and 4.8% of all vaginal samples and 89.6% and 6.2 % in follicular fluid) (Figure 2A and Supplementary figure 2). Indeed, *L. iners* ASV1 was the dominant ASV in 37 out of 112 female samples (33%) (Supplementary figure 2). As expected, *L. crispatus* was also highly prevalent in female samples, being the dominant species in 20 female samples (8 follicular fluid samples and 12 vaginal samples). Generally, despite a similar prevalence between the two types of samples, lactobacilli were less abundant in the follicular fluid compared to vaginal samples. Other highly prevalent species in female samples were *Gardnerella vaginalis* (found in 70% of vaginal samples and 31% follicular fluid samples) and *Prevotella bivia* (found in 62% of vaginal samples and 21% of follicular fluid samples). In male samples, *L. iners* (ASV1) was also the most prevalent bacterium, although present at low estimated abundance (10 out of 92 male samples, 11%).

**Figure 2.**
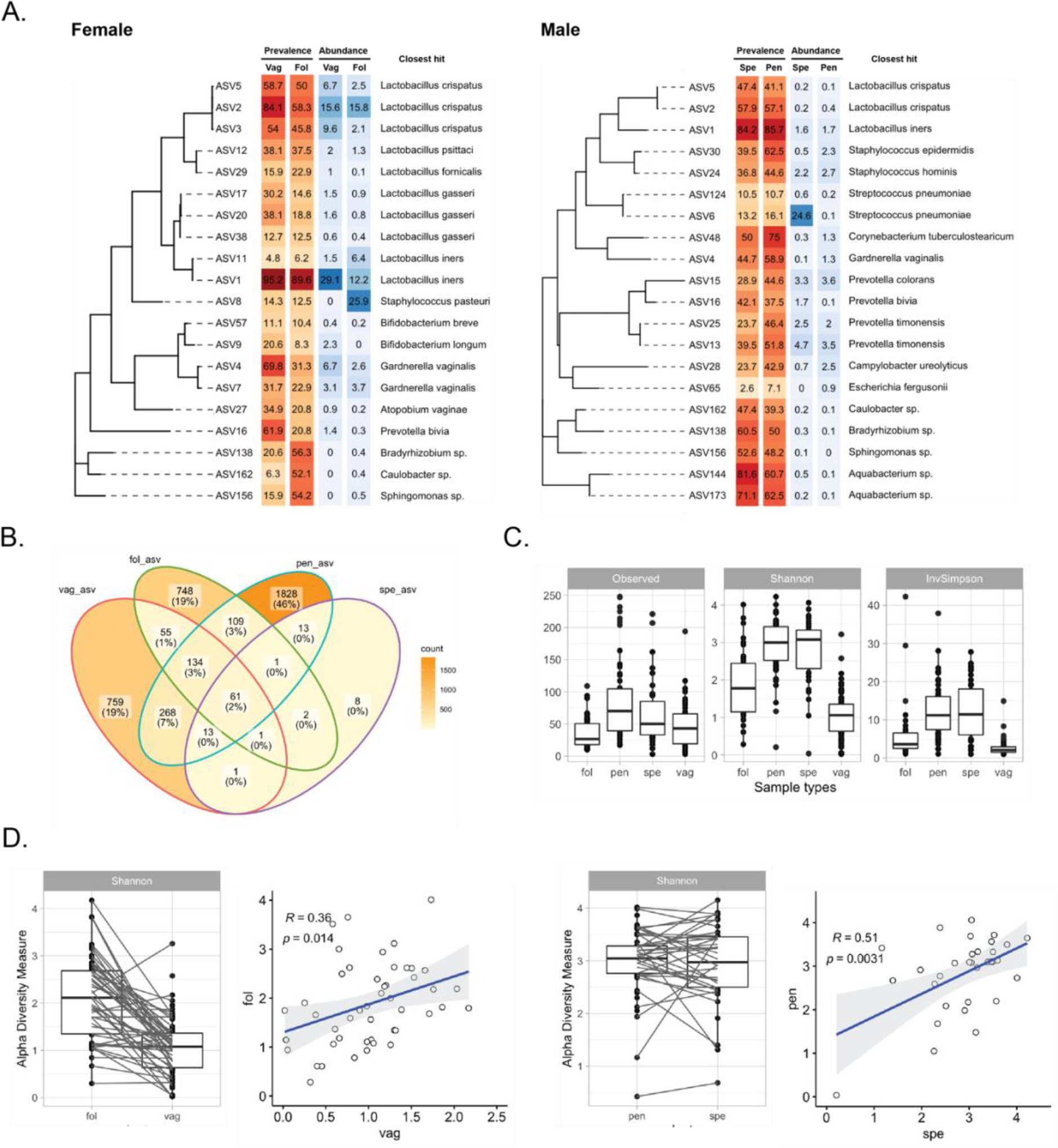
A. Most prevalent ASVs in the female samples and male samples. Sequence alignment generated of 20 most prevalent ASVs using SINA and phylogenetic tree made using FastTree (left panel). Prevalence (red heatmap) and estimated abundance (blue heatmap) of each ASV are shown as percentages. The closest hit (100% coverage and 99% sequence identity) is shown for each ASV on the right panel. B. Shared ASVs between sample types. Venn diagram of shared ASVs across the four sample types. C. Number of observed ASVs, Shannon and inverse Simpson indexes of the four sample types. D. Correlation of Shannon index between samples from same individuals. Boxplot of Shannon indexes of vaginal-follicular fluid (left panel) and sperm-penis (right panel) samples, in which samples from the same individual are connected by the black lines. On the right correlation plots between the samples, with the correlation coefficient and statistical significance. The regression line and the confidence interval are depicted in blue and grey, respectively.

Other prevalent bacteria included previously described genital bacteria like *Gardnerella vaginalis* and multiple *Prevotella* species. Skin-associated bacteria like *Corynebacterium spp.* and *Staphylococcus epidermidis* were also detected, implying their presence on the external male genitalia. In both female and male samples, a cluster Alphaproteobacteria (*Aquabacterium*, *Bradyrhizobium, Sphingomonas and Caulobacter spp.*) may represent putative contaminants that have not been eliminated by our filtering steps.

Since samples were obtained from couples who were presumably sexually active, we wanted to investigate whether any of the genital niches included in the study shared bacterial species. We observed that only 1.52% (n=61) of all ASVs were found in all four sample types were shared amongst these sites (Figure 2B), indicating the resilience of individual site-specific microbial consortia. The main shared genera were *Lactobacillus spp.* (n=21), *Prevotella spp.* (n=10*), Staphylococcus spp.* (n=8) and *Ezakiella spp.* (n= 7).

Further, to characterise the bacterial community structure and species diversity of the four genital niches, we performed alpha diversity analysis (observed number of ASVs, Shannon index and inverted Simpson index) (Figure 2C). As expected, vaginal samples showed the lowest bacterial diversity and evenness (lowest Shannon index and Inverted Simpson index, respectively), as most of them were dominated by *Lactobacillus spp.* Male genital samples showed the highest alpha diversity indices. While for penis samples this may represent a true species variability, given the global low bacterial biomass of this sample group, it may not be true for sperm samples. We did not observe any significant association of alpha diversity metrics and clinical data when analysing sample types separately or combined, except for follicular fluid samples and recurrent implantation failure (Supplementary table 1).

As the lower genital tract may influence the bacterial composition of the upper genital tract, we calculated the Pearson correlation coefficient for intra-individual sample pairs (vagina vs follicular fluid and penis vs sperm – Figure 2D). A moderate positive but significant correlation was found between Shannon indexes of penis and sperm samples (R^2^ = 0.51, P= 0.0031), while a weak positive but significant correlation was obtained for female samples (R^2^ = 0.36, P= 0.014).

To quantify the similarities between bacterial communities and discover differences between sample types, we performed beta diversity analysis using both unweighted and weighted Unifrac distances (Figure 3). These results showed that the four sample types were indeed different from each other and these differences become more apparent when partially overlap when only phylogenetic distance compared (unweighted Unifrac distance) was compared (Figure 3A). Nevertheless, pairwise comparison of beta diversity showed that all sample pairs were significantly different. The same was true when both phylogeny and estimated abundance were considered (weighted Unifrac) (Figure 3B). Here, most of the vaginal samples clustered together, given the high *Lactobacillus spp.* dominance.

**Figure 3.**
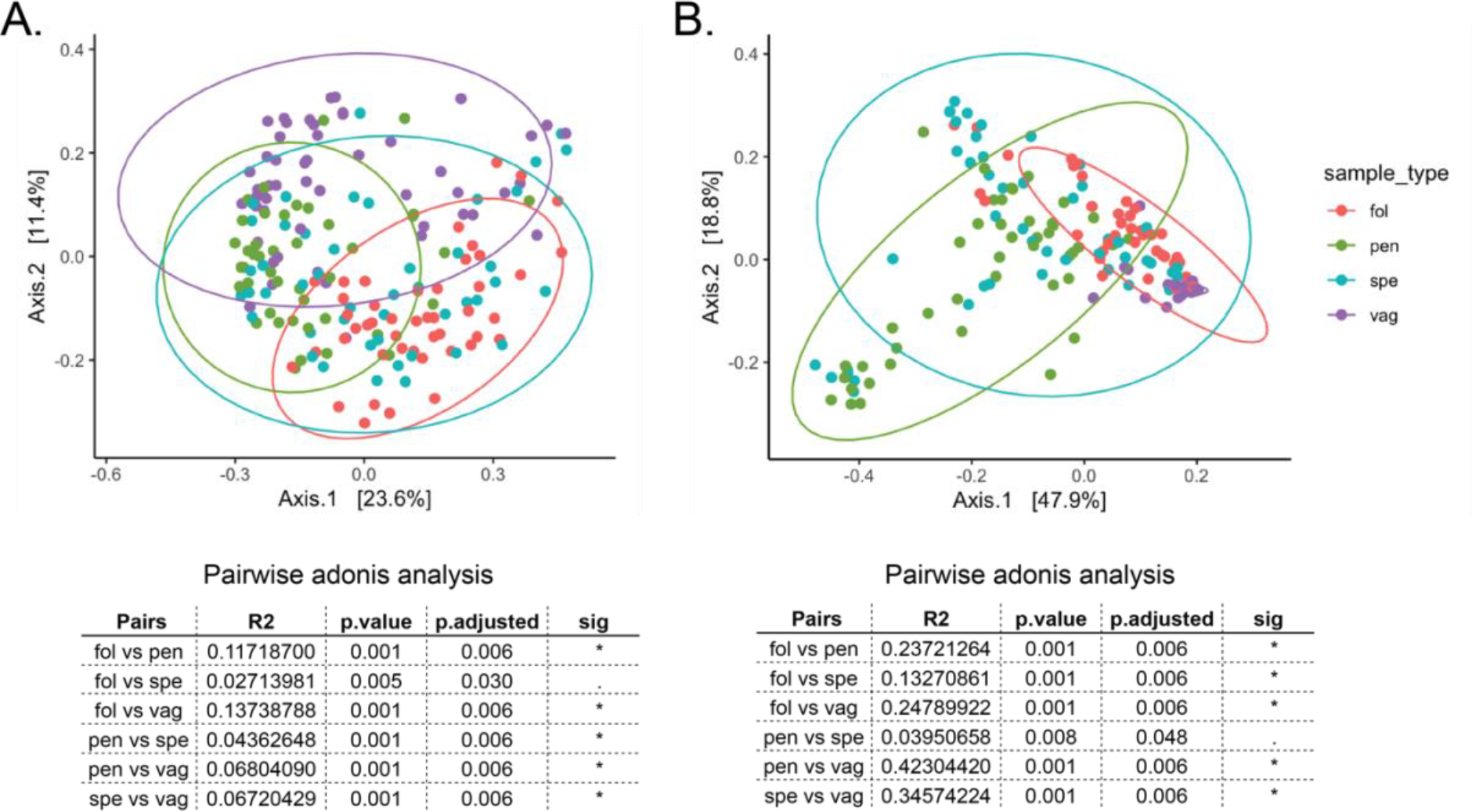
Principal component analysis of (A) unweighted Unifrac and (B) weighted Unifrac distances. Lower panels show permutational multivariate analysis of variance between sample type pairs. Significant values (sig): p<0.01 *, p<0.05.

### Analysis across individuals highlight potential interactions between genital microbiota

To estimate whether there were any microbiota interactions between partners, we compared the beta diversity between samples from different individuals and samples from the same couple (Figure 4A). Despite generally high values, in several cases intra-couple dissimilarities were significantly lower when compared to intra-sample values (same sample type of different individuals). Semen and penis samples from the same individual were more similar compared to semen or penis samples from other individuals. Interestingly, the same was observed for follicular fluid and sperm samples, although this may be the caused by the presence of remaining contaminant ASVs present in these low bacterial biomass samples. On the other hand, intra-individual dissimilarities of vaginal samples were lower compared to other samples.

**Figure 4.**
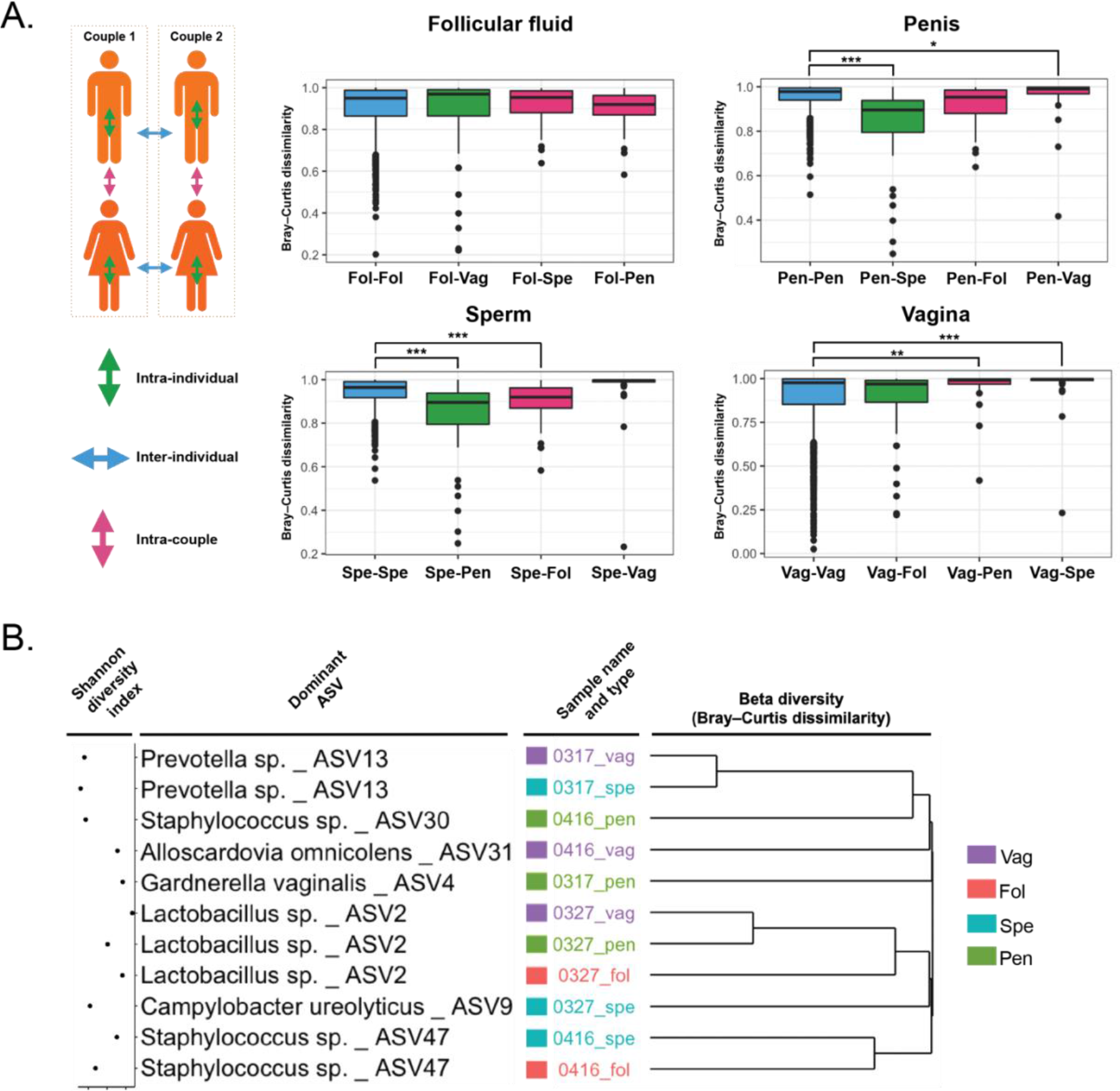
A. Comparison of intra- and inter-sample dissimilarities. The left panel depicts the schematic representation of sample comparisons (intra-individual, inter-individuals and intra-couple). On the right, Bray-Curtis dissimilarities calculated for each sample type. B. Hierarchical clustering dendrogram from the Bray-Curtis dissimilarity analysis in couples in which bacterial transmission was suspected. For each sample, the dominant ASV with species identification is displayed, along with the Shannon diversity index. Statistical analysis was performed by one-way analysis of variance (ANOVA) followed by Tukey’s post hoc test. p<0.01 *, p<0.001 **, p<0.0001 ***.

To further explore inter-sample relationships from the same couple, Bray-Curtis dissimilarities were used to cluster samples based on their community composition (Supplementary figure 3). On several occasions sample pairs from the same patient clustered together, with variable dissimilarity values. This occurrence was higher in male samples (31% of patients in which both sperm and penis samples were present) compared to female samples (13%). Despite the possibility of contamination, this may suggest a possible influence of genital sites on each other. More specifically, given their higher bacterial load, vagina and penis may influence the bacterial composition of follicular fluid and sperm, respectively.

A similar analysis was carried out in couples where inter-partner microbiota interaction was suspected (Figure 4B). The lowest dissimilarity values were observed for vaginal and penis samples from couple 0327 and between vaginal and sperm samples from couple 0317. In the first case, the dominant ASV in both sample types was *Lactobacillus sp.* ASV2. While lactobacilli are not frequently found in penis samples, this may suggest a transfer of microbiota from the vagina (*Lactobacillus sp.* ASV2 is also found in the follicular sample) to the penis. In the second case, an influence of the male microbiota on the female colonisation could be suspected as the dominant ASV was *Prevotella sp.* ASV13 in both samples. This is further highlighted by the presence of two additional *Prevotella spp.* (ASV25 and ASV15) as the second and third dominant ASVs in both sample types.

### Specifically enriched genera

Next, we sought to identify differentially abundant taxa (at genus level) specifically enriched between sites in female and male samples (Figure 5).

**Figure 5.**
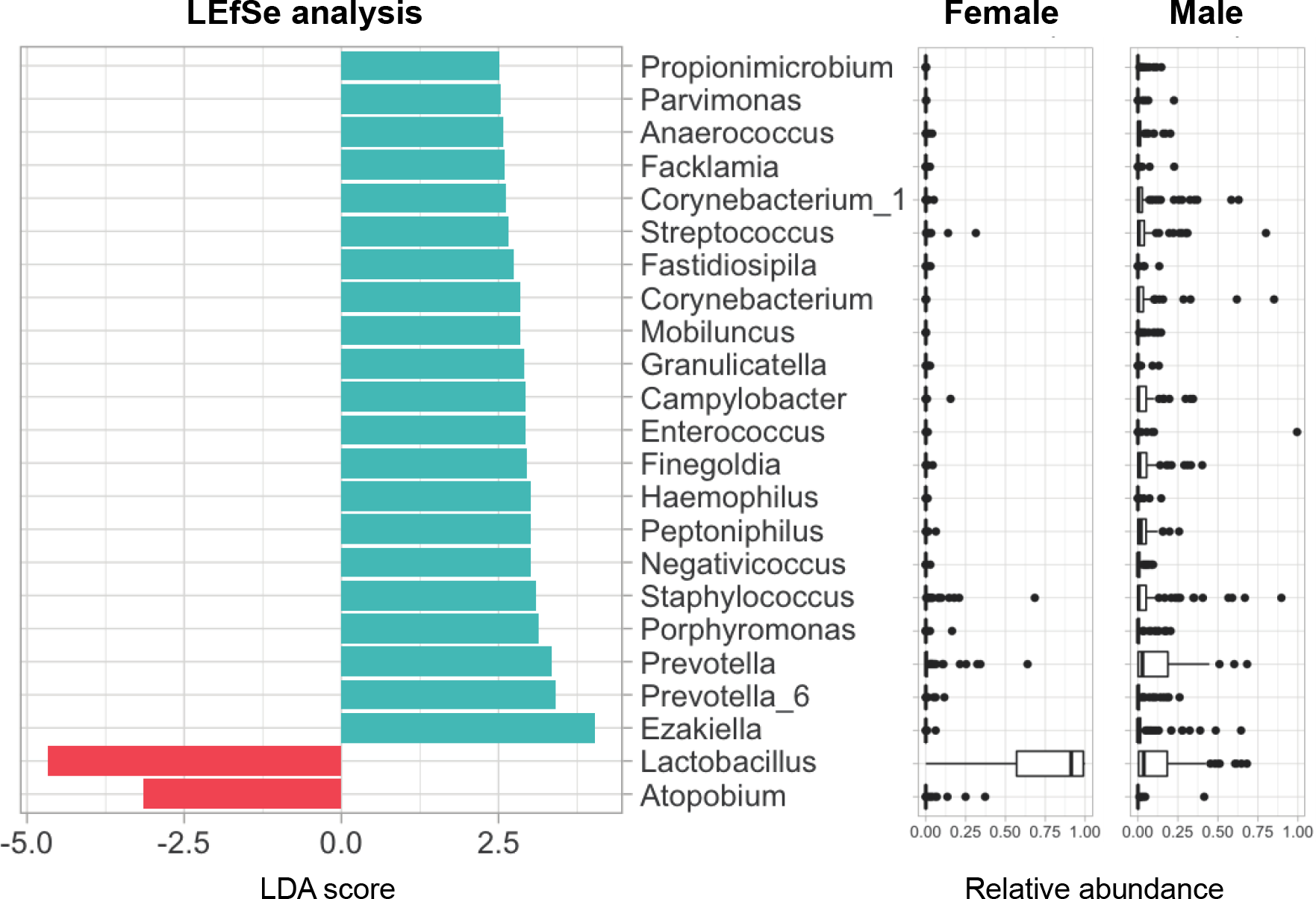
Signature genera in female and male samples. Barplots depicting the effect size (log10 transformed LDA score) for specific genera. Relative estimated abundance of depicted genera is displayed on the right panels for female and male samples.

As expected, genus *Lactobacillus* was enriched in female samples, along with the genus *Atopobium*. In male samples, a number of genera were specifically enriched, the majority of which were previously associated with negative gynecological and obstetrical outcomes. These included *Prevotella*, *Porphyromonas*, *Peptoniphilus*, *Finegoldia*, *Campylobacter*, *Mobiluncus* among others.

## Discussion

Despite the progress in the human reproduction medicine, an increasing number of infertility cases are still considered as idiopathic. Reproductive success may be influenced by multiple causes, including genetic, physiological and environmental causes. Since bacteria have an increasingly recognised influence on human homeostasis, it is tempting to explore the influence of the microbiota on human reproduction. Previous studies highlighted the impact of vaginal microbiota on pregnancy and sexually transmitted diseases [48–51]. Nevertheless, there is a knowledge gap in the field of genital microbiota compared to the plethora of studies about the influence of bacteria on other body sites. Most of the studies focus on the vagina, while the microbiota of the upper female and the male genital tract remain overlooked. Moreover, it is still unknown on how sexual activity may influence the bacterial colonisation of female genital tract and impact the fate of pregnancy.

In this study, we explored the composition of genital microbiota in infertile couples, which included samples from both female and male partners. Given the limitations in the sampling of the upper genital, which may be highly invasive, we used samples that are routinely obtained during ART procedures, namely the follicular fluid (obtained during oocytes retrieval) and semen (used for the spermiogram analysis and *in vitro* fertilisation). Sampling of the lower genital tract included vaginal and penile swabs.

Bacterial biomass was highly variable between sample types varying according to the sampling site on the body. As expected, *Lactobacillus spp.* were highly prevalent in vaginal samples, while the penis glans samples were moderately colonised by multiple bacterial species. On the other hand, follicular fluid and semen samples had generally low bacterial biomass. Here we also show unlike the vagina, other sampled body sites (follicular fluids, penis and semen) were primarily of low bacterial biomass. With increasing emphasis on proper reporting of microbial species from multiple human body sites, we applied stringent filtering procedure to limit the effect of possible contaminations on the results and use absolute bacterial copy numbers while reporting.

As expected, *Lactobacillus* was the dominant genus among female samples, with a relative abundance above 90% in 45 out of 63 vaginal samples. Moreover, it was the dominant genus in 39 out of 48 follicular fluid samples included in the analysis. Previous studies suggested that vaginal microbiota may influence the colonisation of the upper genital tract. Nevertheless, we observed a related microbiota pattern in only 13% of women in which both vaginal and follicular fluid samples were present, while the remaining samples displayed differences in colonisation pattern and the dominant species. Thus, it appears that the upper female genital tract may have a distinctive low biomass microbiota, as previously suggested [15,52,53]. Further studies are required to characterise this specific microbiota, which is challenging due to the invasiveness of sampling and high risk of contamination, principally by the vaginal bacteria.

Male samples were highly diverse compared to female samples. Despite this, semen and penis samples from the same individual, were more similar compared to samples of the same types from other men. This suggests that the penile microbiota and its relatively high bacterial load, may have an important influence on the bacterial composition of the ejaculate. Our results showed that male samples were specifically enriched with different bacterial genera previously associated with bacterial vaginosis (*Prevotella*, *Porphyromonas*, *Peptoniphilus*, *Finegoldia*, *Campylobacter* and *Mobiluncus* genera, among others) and therefore may represent a potential reservoir.

Previous reports have shown that sexual intercourses may have an influence on the bacterial colonisation of the vagina, with a decrease in the relative abundance of *Lactobacillus sp.* [36,54,55]. We highlight here a possible inter-partner interaction of microbiota in two cases. In the first one, penile microbiota composition was very similar to the partner’s vaginal microbiota, with *Lactobacillus sp.* ASV2 being the dominant ASV in both sample types, thus indicating a female to male bacterial transmission. In the second case, microbiota composition of the semen was highly similar to the partner’s vaginal microbiota. In this case, *Prevotella sp.* ASV13 was the dominant bacterium in both samples, thus suggesting a male to female transmission. Nevertheless, the lack of information about the sexual activity of the couples included in the study is an important limitation for drawing strong conclusions on bacterial transmission between partners.

Future analysis of the influence of sexual activity on microbial composition and variation of genital sites should ideally comprise samples obtained before and after sexual intercourse. Moreover, additional studies about the stability of the seminal microbiota based on longitudinal sampling of both penile swabs and semen should be performed.

## References

1. Valdes AM, Walter J, Segal E, Spector TD. Role of the gut microbiota in nutrition and health. BMJ [Internet]. 2018 [cited 2020 Aug 30];361. Available from: https://www.bmj.com/content/361/bmj.k2179

2. Gilbert JA, Blaser MJ, Caporaso JG, Jansson JK, Lynch SV, Knight R. Current understanding of the human microbiome. Nature Medicine. 2018;24:392–400.

3. Dominguez-Bello MG, Godoy-Vitorino F, Knight R, Blaser MJ. Role of the microbiome in human development. Gut. 2019;68:1108–14.

4. Robertson RC, Manges AR, Finlay BB, Prendergast AJ. The Human Microbiome and Child Growth – First 1000 Days and Beyond. Trends in Microbiology. 2019;27:131–47.

5. Gupta P, Singh MP, Goyal K. Diversity of Vaginal Microbiome in Pregnancy: Deciphering the Obscurity. Front Public Health. 2020;8:326.

6. O’Callaghan JL, Turner R, Dekker Nitert M, Barrett HL, Clifton V, Pelzer ES. Re-assessing microbiomes in the low-biomass reproductive niche. BJOG. 2020;127:147–58.

7. Heil BA, Paccamonti DL, Sones JL. Role for the mammalian female reproductive tract microbiome in pregnancy outcomes. Physiol Genomics. 2019;51:390–9.

8. France M, Alizadeh M, Brown S, Ma B, Ravel J. Towards a deeper understanding of the vaginal microbiota. Nat Microbiol. 2022;7:367–78.

9. Gajer P, Brotman RM, Bai G, Sakamoto J, Schutte UME, Zhong X, et al. Temporal Dynamics of the Human Vaginal Microbiota. Science Translational Medicine. 2012;4:132ra52- 132ra52.

10. Hickey RJ, Zhou X, Settles ML, Erb J, Malone K, Hansmann MA, et al. Vaginal microbiota of adolescent girls prior to the onset of menarche resemble those of reproductive-age women. mBio. 2015;6:e00097–15.

11. Muhleisen AL, Herbst-Kralovetz MM. Menopause and the vaginal microbiome. Maturitas. 2016;91:42–50.

12. Ravel J, Gajer P, Abdo Z, Schneider GM, Koenig SSK, McCulle SL, et al. Vaginal microbiome of reproductive-age women. Proc Natl Acad Sci USA. 2011;108 Suppl 1:4680–7.

13. Mehta SD, Nannini DR, Otieno F, Green SJ, Agingu W, Landay A, et al. Host Genetic Factors Associated with Vaginal Microbiome Composition in Kenyan Women. mSystems. 2020;5:e00502–20.

14. Peric A, Weiss J, Vulliemoz N, Baud D, Stojanov M. Bacterial Colonization of the Female Upper Genital Tract. International Journal of Molecular Sciences. 2019;20:3405.

15. Chen C, Song X, Wei W, Zhong H, Dai J, Lan Z, et al. The microbiota continuum along the female reproductive tract and its relation to uterine-related diseases. Nature Communications [Internet]. 2017 [cited 2017 Nov 28];8. Available from: http://www.nature.com/articles/s41467-017-00901-0

16. Moreno I, Franasiak JM. Endometrial microbiota—new player in town. Fertility and Sterility. 2017;108:32–9.

17. Onderdonk AB, Delaney ML, Fichorova RN. The Human Microbiome during Bacterial Vaginosis. Clinical Microbiology Reviews. 2016;29:223–38.

18. Schellenberg JJ, Patterson MH, Hill JE. Gardnerella vaginalis diversity and ecology in relation to vaginal symptoms. Research in Microbiology. 2017;168:837–44.

19. Ravel J, Brotman RM, Gajer P, Ma B, Nandy M, Fadrosh DW, et al. Daily temporal dynamics of vaginal microbiota before, during and after episodes of bacterial vaginosis. Microbiome. 2013;1:29.

20. Tang W, Mao J, Li KT, Walker JS, Chou R, Fu R, et al. Pregnancy and fertility-related adverse outcomes associated with Chlamydia trachomatis infection: a global systematic review and meta-analysis. Sex Transm Infect. 2020;96:322–9.

21. Jung H-S, Ehlers MM, Lombaard H, Redelinghuys MJ, Kock MM. Etiology of bacterial vaginosis and polymicrobial biofilm formation. Critical Reviews in Microbiology. 2017;43:651 – 67.

22. van de Wijgert JHHM, Borgdorff H, Verhelst R, Crucitti T, Francis S, Verstraelen H, et al. The Vaginal Microbiota: What Have We Learned after a Decade of Molecular Characterization? Fredricks DN, editor. PLoS ONE. 2014;9:e105998.

23. Smith SB, Ravel J. The vaginal microbiota, host defence and reproductive physiology. J Physiol (Lond). 2017;595:451–63.

24. Amabebe E, Anumba DOC. The Vaginal Microenvironment: The Physiologic Role of Lactobacilli. Frontiers in Medicine [Internet]. 2018 [cited 2018 Oct 5];5. Available from: https://www.frontiersin.org/article/10.3389/fmed.2018.00181/full

25. Satta A. Experimental Chlamydia trachomatis infection causes apoptosis in human sperm. Human Reproduction. 2005;21:134–7.

26. Wølner-Hanssen P, Mårdh P-A. In vitro tests of the adherence of Chlamydia trachomatis to human spermatozoa. Fertility and Sterility. 1984;42:102–7.

27. Nunez-Calonge R, Caballero P, Redondo C, Baquero F, Martinez-Ferrer M, Meseguer MA. Ureaplasma urealyticum reduces motility and induces membrane alterations in human spermatozoa. Human Reproduction. 1998;13:2756–61.

28. Ahmadi MH, Mirsalehian A, Sadighi Gilani MA, Bahador A, Talebi M. Asymptomatic Infection With Mycoplasma hominis Negatively Affects Semen Parameters and Leads to Male Infertility as Confirmed by Improved Semen Parameters After Antibiotic Treatment. Urology. 2017;100:97–102.

29. Baud D, Pattaroni C, Vulliemoz N, Castella V, Marsland BJ, Stojanov M. Sperm Microbiota and Its Impact on Semen Parameters. Frontiers in Microbiology [Internet]. 2019 [cited 2019 Mar 8];10. Available from: https://www.frontiersin.org/article/10.3389/fmicb.2019.00234/full

30. Hou D, Zhou X, Zhong X, Settles ML, Herring J, Wang L, et al. Microbiota of the seminal fluid from healthy and infertile men. Fertility and Sterility. 2013;100:1261–1269.e3.

31. Weng S-L, Chiu C-M, Lin F-M, Huang W-C, Liang C, Yang T, et al. Bacterial Communities in Semen from Men of Infertile Couples: Metagenomic Sequencing Reveals Relationships of Seminal Microbiota to Semen Quality. PLOS ONE. 2014;9:e110152.

32. Štšepetova J, Baranova J, Simm J, Parm Ü, Rööp T, Sokmann S, et al. The complex microbiome from native semen to embryo culture environment in human in vitro fertilization procedure. Reprod Biol Endocrinol. 2020;18:3.

33. Alfano M, Ferrarese R, Locatelli I, Ventimiglia E, Ippolito S, Gallina P, et al. Testicular microbiome in azoospermic men-first evidence of the impact of an altered microenvironment. Hum Reprod. 2018;33:1212–7.

34. Koedooder R, Singer M, Schoenmakers S, Savelkoul PHM, Morré SA, de Jonge JD, et al. The vaginal microbiome as a predictor for outcome of in vitro fertilization with or without intracytoplasmic sperm injection: a prospective study. Hum Reprod. 2019;34:1042–54.

35. Koedooder R, Singer M, Schoenmakers S, Savelkoul PHM, Morré SA, de Jonge JD, et al. The ReceptIVFity cohort study protocol to validate the urogenital microbiome as predictor for IVF or IVF/ICSI outcome. Reprod Health. 2018;15:202.

36. Mändar R, Punab M, Borovkova N, Lapp E, Kiiker R, Korrovits P, et al. Complementary seminovaginal microbiome in couples. Research in Microbiology. 2015;166:440–7.

37. Borovkova N, Korrovits P, Ausmees K, Türk S, Jõers K, Punab M, et al. Influence of sexual intercourse on genital tract microbiota in infertile couples. Anaerobe. 2011;17:414–8.

38. Rapin A, Pattaroni C, Marsland BJ, Harris NL. Microbiota Analysis Using an Illumina MiSeq Platform to Sequence 16S rRNA Genes. Curr Protoc Mouse Biol. 2017;7:100–29.

39. Eren AM, Vineis JH, Morrison HG, Sogin ML. A Filtering Method to Generate High Quality Short Reads Using Illumina Paired-End Technology. PLOS ONE. 2013;8:e66643.

40. Callahan BJ, McMurdie PJ, Rosen MJ, Han AW, Johnson AJA, Holmes SP. DADA2: High resolution sample inference from Illumina amplicon data. Nat Methods. 2016;13:581–3.

41. Quast C, Pruesse E, Yilmaz P, Gerken J, Schweer T, Yarza P, et al. The SILVA ribosomal RNA gene database project: improved data processing and web-based tools. Nucleic Acids Res. 2013;41:D590–596.

42. Davis NM, Proctor DM, Holmes SP, Relman DA, Callahan BJ. Simple statistical identification and removal of contaminant sequences in marker-gene and metagenomics data. Microbiome. 2018;6:226.

43. McMurdie PJ, Holmes S. phyloseq: An R Package for Reproducible Interactive Analysis and Graphics of Microbiome Census Data. PLOS ONE. 2013;8:e61217.

44. Lahti L, Shetty S. “microbiome R package.” 2022;

45. Jari Oksanen, Gavin L. Simpson, F. Guillaume Blanchet, Roeland Kindt, Pierre Legendre, Peter R. Minchin, R.B., O’Hara, Peter Solymos, M. Henry H. Stevens, Eduard Szoecs, Helene Wagner, Matt Barbour, Michael Bedward, Ben, Bolker, Daniel Borcard, Gustavo Carvalho, Michael Chirico, Miquel De Caceres, Sebastien Durand, Heloisa Beatriz, Antoniazi Evangelista, Rich FitzJohn, Michael Friendly, Brendan Furneaux, Geoffrey Hannigan, Mark O. Hill, Leo, Lahti, Dan McGlinn, Marie-Helene Ouellette, Eduardo Ribeiro Cunha, Tyler Smith, Adrian Stier, Cajo J.F. Ter Braak, and James Weedon. vegan: Community Ecology Package. 2022;

47. Asya Khleborodova. lefser: R implementation of the LEfSE method for microbiome biomarker discovery. R package version 1.2.0. https://github.com/waldronlab/lefser. 2021;

47. Molina NM, Plaza-Díaz J, Vilchez-Vargas R, Sola-Leyva A, Vargas E, Mendoza-Tesarik R, et al. Assessing the testicular sperm microbiome: a low-biomass site with abundant contamination. Reproductive BioMedicine Online. 2021;43:523–31.

48. Tabatabaei N, Eren A, Barreiro L, Yotova V, Dumaine A, Allard C, et al. Vaginal microbiome in early pregnancy and subsequent risk of spontaneous preterm birth: a case - control study. BJOG: An International Journal of Obstetrics & Gynaecology. 2019;126:349– 58.

49. Fettweis JM, Serrano MG, Brooks JP, Edwards DJ, Girerd PH, Parikh HI, et al. The vaginal microbiome and preterm birth. Nature Medicine. 2019;25:1012–21.

50. Nelson DB, Rockwell LC, Prioleau MD, Goetzl L. The role of the bacterial microbiota on reproductive and pregnancy health. Anaerobe. 2016;42:67–73.

51. Green KA, Zarek SM, Catherino WH. Gynecologic health and disease in relation to the microbiome of the female reproductive tract. Fertility and Sterility. 2015;104:1351–7.

52. Li F, Chen C, Wei W, Wang Z, Dai J, Hao L, et al. The metagenome of the female upper reproductive tract. GigaScience [Internet]. 2018 [cited 2019 Mar 14];7. Available from: https://academic.oup.com/gigascience/article/doi/10.1093/gigascience/giy107/5091799

53. Mitchell CM, Haick A, Nkwopara E, Garcia R, Rendi M, Agnew K, et al. Colonization of the upper genital tract by vaginal bacterial species in nonpregnant women. American Journal of Obstetrics and Gynecology. 2015;212:611.e1–611.e9.

54. Vodstrcil LA, Twin J, Garland SM, Fairley CK, Hocking JS, Law MG, et al. The influence of sexual activity on the vaginal microbiota and Gardnerella vaginalis clade diversity in young women. Fredricks DN, editor. PLOS ONE. 2017;12:e0171856.

55. Carter KA, France MT, Rutt L, Bilski L, Martinez-Greiwe S, Regan M, et al. Sexual transmission of urogenital bacteria: whole metagenome sequencing evidence from a sexual network study. mSphere. 2024;9:e00030–24.

