## Supplementary figure 1 for "Genital microbiota in infertile couples"

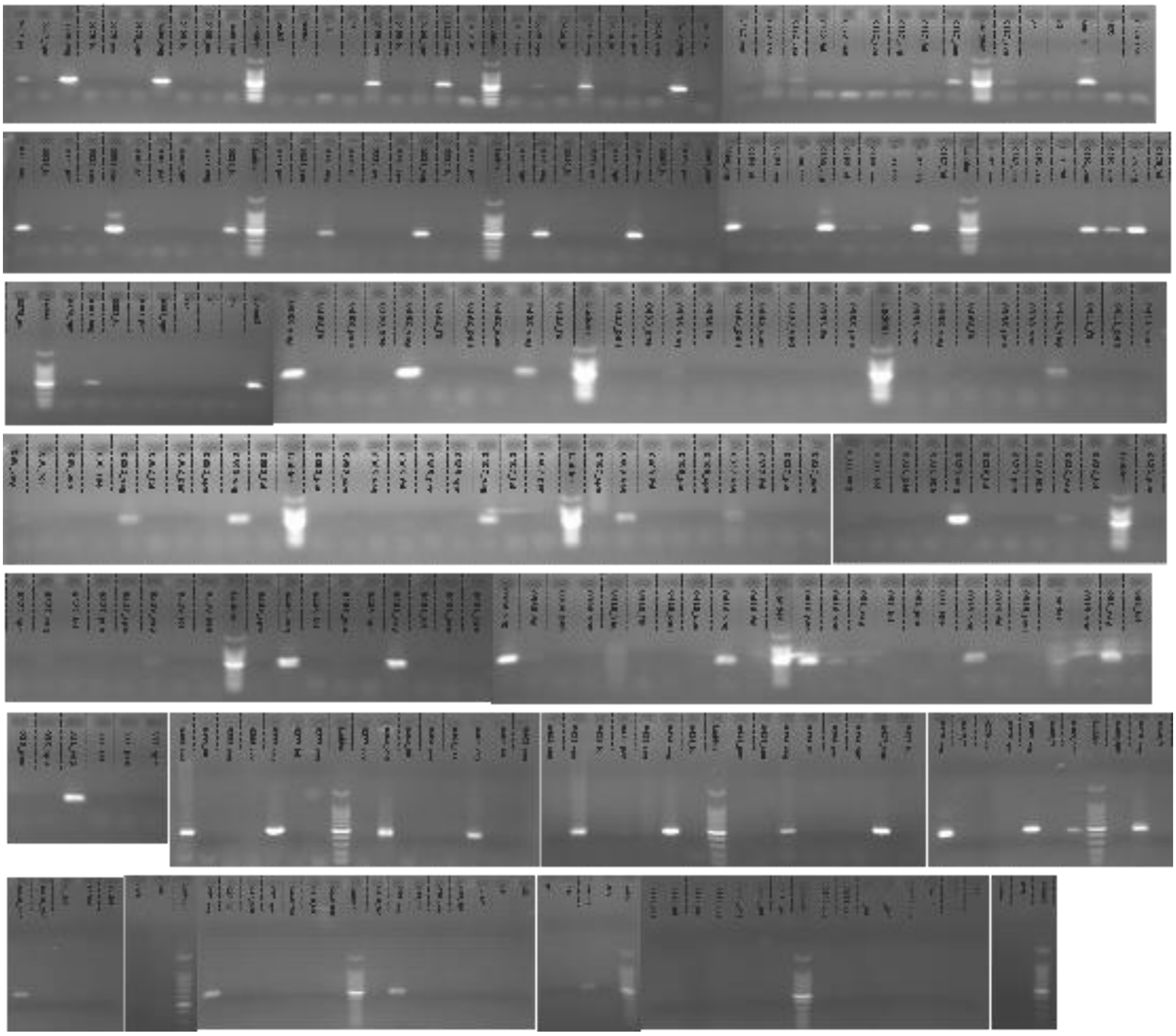

Supplementary figure 1 – Evaluation of the PCR product after the amplification of the V1-V2 region of the 16S rRNA gene. Samples were categorized based on band intensity and used for sequencing libraries preparation.
