## Supplementary figure 2 for "Genital microbiota in infertile couples"

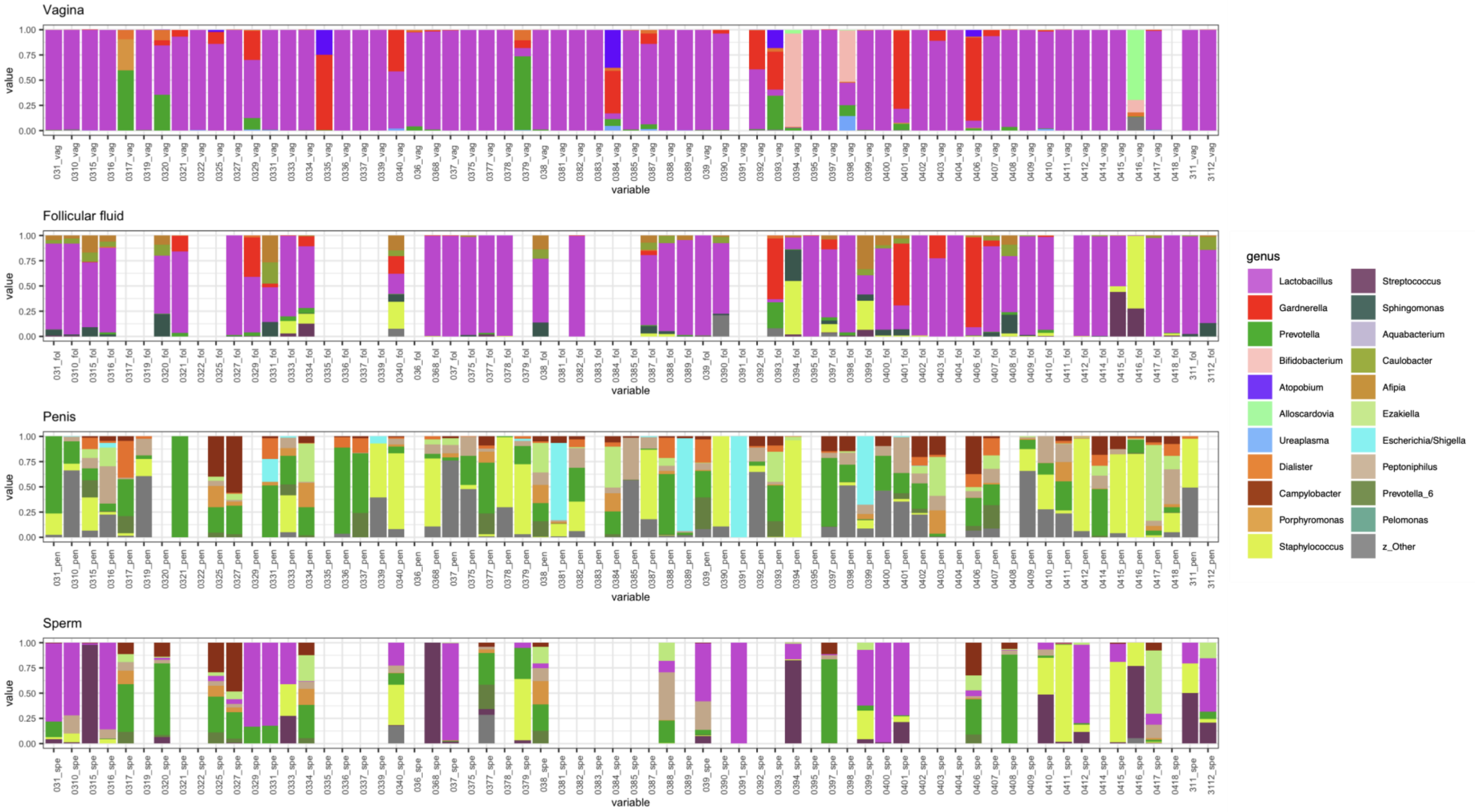


Supplementary figure 2 – Percent stacked barplot of the most abundant genera in the four sample types. Samples belonging to the same couple are aligned vertically. Missing bars indicate samples that did not pass the filtration steps indicated in the manuscript.
