## Supplementary figure 3 for "Genital microbiota in infertile couples"

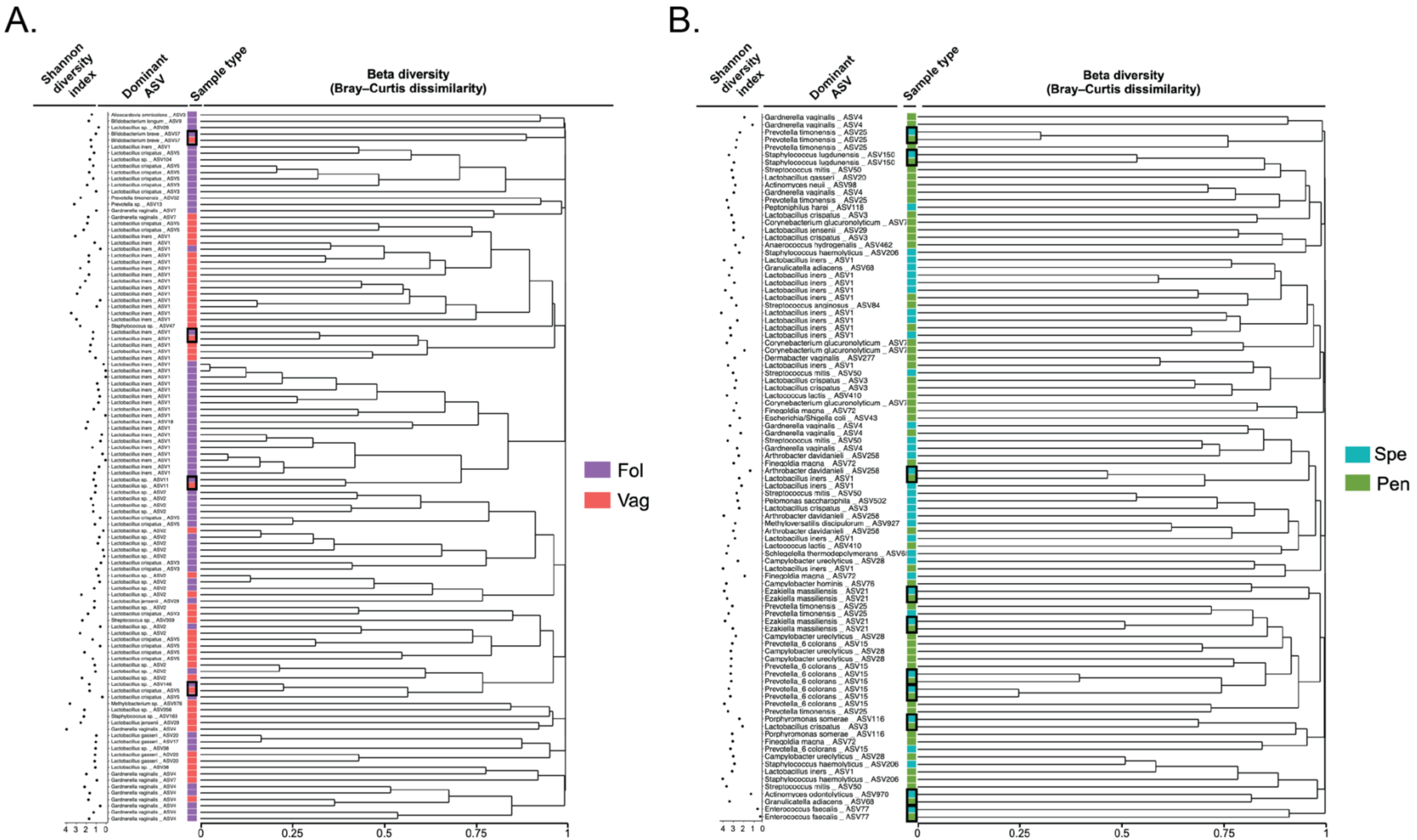


Supplementary figure 3 – Clustering of the samples based on community composition. Hierarchical clustering dendrogram from the Bray-Curtis dissimilarity analysis of (A) female and (B) male samples based on Bray-Curtis dissimilarity (right panel). For each sample, the dominant ASV with species identification is displayed, along with the Shannon diversity index. Black boxes depict samples from the same patient.
