## Supplementary table 1 for "Genital microbiota in infertile couples"

Supplementary table 1 – Association of alpha diversity metrics and clinical variables

| Clinical variables | Average Shannon index "yes" group | Average Shannon index "no" group | T statistics | P-value |
|---|---|---|---|---|
| **Female samples** | | | | |
| Miscarriage | 1.59 | 1.41 | 0.67 | 0.51 |
| Tubal factor | 1.49 | 1.40 | 0.53 | 0.60 |
| Recurrent implantation failure | 1.51 | 1.43 | 0.23 | 0.83 |
| Previous miscarriage | 1.38 | 1.45 | -0.32 | 0.75 |
| Recurrent miscarriage | 0.05 | 1.45 | -0.32 | 0.75 |
| Endometriosis | 1.49 | 1.42 | 0.33 | 0.74 |
| Myometritis | 1.80 | 1.40 | 1.44 | 0.18 |
| **Follicular fluid samples** | | | | |
| Miscarriage | 2.33 | 1.84 | 1.58 | 0.16 |
| Tubal factor | 1.97 | 1.86 | 0.54 | 0.59 |
| Recurrent implantation failure | 2.32 | 1.88 | 3.01 | 0.01 |
| Previous miscarriage | 1.83 | 1.91 | -0.31 | 0.76 |
| Recurrent miscarriages | NA | 1.90 | NA | NA |
| Endometriosis | 2.00 | 1.87 | 0.43 | 0.67 |
| Myometritis | 2.49 | 1.83 | 2.32 | 0.06 |
| **Vaginal samples** | | | | |
| Miscarriage | 0.96 | 1.09 | -0.54 | 0.60 |
| Tubal factor | 1.10 | 1.06 | 0.22 | 0.83 |
| Recurrent implantation failure | 0.97 | 1.08 | -0.44 | 0.69 |
| Previous miscarriage | 1.08 | 1.07 | 0.01 | 0.99 |
| Recurrent miscarriage | 0.05 | 1.09 | 0.01 | 0.99 |
| Endometriosis | 1.02 | 1.09 | -0.32 | 0.75 |
| Myometritis | 1.11 | 1.07 | 0.27 | 0.79 |
| **Male samples** | | | | |
| Normal spermiogram | 2.89 | 2.80 | 0.62 | 0.54 |
| Oligozoospermia | 2.94 | 2.78 | 1.00 | 0.32 |
| Asthenozoospermia | 2.83 | 2.83 | -0.03 | 0.98 |
| Teratozoospermia | 2.79 | 2.86 | -0.43 | 0.67 |
| Azoospermia | 2.12 | 2.85 | -0.85 | 0.55 |
| Miscarriage | 2.98 | 2.81 | 0.77 | 0.46 |
| Recurrent miscarriages | NA | 2.83 | NA | 0.46 |
| **Penis samples** | | | | |
| Normal spermiogram | 2.88 | 2.81 | 0.45 | 0.65 |
| Oligozoospermia | 2.85 | 2.84 | 0.05 | 0.96 |
| Asthenozoospermia | 2.79 | 2.86 | -0.33 | 0.75 |
| Teratozoospermia | 2.80 | 2.86 | -0.29 | 0.77 |
| Azoospermia | 2.97 | 2.84 | -0.29 | 0.77 |
| Miscarriage | 3.02 | 2.82 | 0.57 | 0.59 |
| Recurrent miscarriages | NA | 2.84 | NA | 0.59 |
| **Sperm samples** | | | | |
| Normal spermiogram | 2.89 | 2.79 | 0.39 | 0.70 |
| Oligozoospermia | 3.07 | 2.69 | 1.39 | 0.18 |
| Asthenozoospermia | 2.87 | 2.78 | 0.31 | 0.76 |
| Teratozoospermia | 2.77 | 2.86 | -0.30 | 0.77 |
| Azoospermia | 1.27 | 2.86 | -0.30 | 0.77 |
| Miscarriage | 2.94 | 2.80 | 0.47 | 0.65 |
| Recurrent miscarriages | NA | 2.82 | NA | 0.65 |
